# Multivariate Pattern Analysis of EEG Reveals Neural Mechanism of Naturalistic Target Processing in Attentional Blink

**DOI:** 10.1101/2023.11.29.569260

**Authors:** Mansoure Jahanian, Marc Joanisse, Boyu Wang, Yalda Mohsenzadeh

## Abstract

The human brain has inherent limitations in consciously processing visual information. When individuals monitor a rapid sequence of images for detecting two targets, they often miss the second target (T2) if it appears within a short time frame of 200-500ms after the first target (T1), a phenomenon known as the attentional blink (AB). The neural mechanism behind AB remains unclear, largely due to the use of simplistic visual items such as letters and digits in conventional AB experiments, which differ significantly from naturalistic vision. This study employed advanced multivariate pattern analysis (MVPA) of human EEG data to explore the neural representations associated with target processing within a naturalistic paradigm under conditions where AB does or does not occur. Our MVPA analysis successfully decoded the identity of target images from EEG data. Moreover, in the AB condition, characterized by a limited time between targets, T1 processing coincided with T2 processing, resulting in the suppression of late representational markers of both T1 and T2. Conversely, in the condition with longer inter-target interval, neural representations endured for a longer duration. These findings suggest that the attentional blink can be attributed to the suppression of neural representations in the later stages of target processing.

**Significance Statement:** Within a naturalistic paradigm, we investigated the phenomenon known as attentional blink, where individuals struggle to identify a second target in a rapid sequence when the first target precedes it too closely. Attentional blink is purported to reflect an apparent bottleneck in the attention system’s ability to rapidly redirect attentional resources; however, the mechanism underlying this phenomenon remains hotly debated. Our findings reveal that during a rapid presentation of natural images, a short temporal gap between targets results in reduced neural representations of targets and the occurrence of attentional blink. Conversely, when a greater temporal gap exists between targets, neural representations are preserved. This study provides valuable insights into how the human brain perceives the ever-changing visual world around us.

## 1 Introduction

In the blink of an eye, the surrounding environment bombards our brain with an influx of visual information, exceeding the brain’s capacity for simultaneous processing. Yet, rather than being overwhelmed, we understand the visual world usually with minimal effort. This feat is possible by visual attention, a mechanism that enables us to perform efficiently in spatial visual searches and pinpoint desired targets among a multitude of distractors (Treisman and Gelade, 1980; Carrasco, 2011). However, with a rapid presentation of images in spatially overlapping locations, it becomes challenging to identify subsequent targets that appear closely in time (Raymond et al., 1992; Potter et al., 2014; Mohsenzadeh et al., 2018, 2019). Raymond et al. (1992) showed that participants fail to correctly report the second of two targets in rapid succession if the second target (T2) occurs within 200 to 500 ms of the first target (T1), a phenomenon which is known as the attentional blink (AB).

In a typical AB paradigm, participants show a decreased performance in reporting T2 under short lag conditions, where the targets are only 200-500 ms apart. Conversely, when the lag is longer, with targets separated by 500 ms or more, T2 reporting performance significantly improves among participants (Raymond et al., 1992; Einhäuser et al., 2007; Tang et al., 2020). Early research suggested that AB is associated with a capacity limitation in the processing of targets, and this limitation prevents T2 from advancing to the later processing stages until T1 processing is completed. (Chun and Potter, 1995; Potter et al., 2002; Dehaene et al., 2003). However, Di Lollo et al. (2005) provided evidence in opposition to the limited capacity theories by showing that participants had no difficulty reporting three consecutive targets chosen from the same category. Therefore, they proposed that AB arises from a temporary loss of control over a specific attentional set. Moreover, recent studies presented evidence supporting that AB arises from a dysfunctional gating of information into working memory rather than a resource limitation (Olivers and Nieuwenhuis, 2006; Olivers and Meeter, 2008). These contradictory findings motivated studies to investigate the temporal aspects of attentional blink; however, the mechanisms involved in this phenomenon have yet to be discovered.

To explore the neural mechanisms of AB with a high temporal resolution, researchers have employed Electroencephalography (EEG) (Zivony and Lamy, 2022). A recent advancement in EEG research is multivariate pattern analysis (MVPA), which can detect subtle differences in the temporal pattern of neural activity that cannot be captured by traditional univariate methods. MVPA can be used to decode the neural representations of target processing based on the temporal patterns (Cichy and Pantazis, 2017). A few recent studies have utilized MVPA to investigate AB, revealing the ability to decode the identities of letter or digit targets through MVPA of EEG/MEG data (Meijs et al., 2019; Alilović et al., 2021). Nonetheless, important questions remain unanswered. It is unclear how neural representations differ when participants are presented with natural stimuli that resemble real-world visual experience. Delving into the study of naturalistic paradigms not only enables us to explore the differences in the AB effect between the natural and traditional (digits and letters) stimuli contexts but also facilitates the expansion of AB models to encompass these intricate stimuli. Additionally, MVPA allows us to isolate neural representations of targets across different lag conditions, providing a fine-grained exploration of AB. Addressing the differences in target processing caused by lag conditions is important given that AB happens in short lag conditions.

Thus, we aimed to explore the neural representations of target processing within an attentional blink paradigm with natural stimuli and compare the representations under short and long lag conditions. To address the aim, we collected EEG and behavioral data while participants viewed rapid series of object images embedded with two face targets, and they were instructed to report targets. We used MVPA of the EEG data to compare the temporal dynamics of T1 and T2 processing in two lag conditions. Our results indicated that attentional blink, observed under short lag conditions, can be linked to the suppression of neural representations for both targets during the later stages of processing.

## 2 Materials and Methods

### 2.1 Participants

Thirty-five healthy adult participants took part in the experiment. One participant was excluded from the analysis due to the chance level performance in the behavior task for a final sample of 34 participants (16 female; age range: 18 - 35 years; mean = 22.9; STD = 3.5). All participants were right-handed and had normal or corrected-to-normal vision. Participants provided informed consent prior to the experiment, and received compensation for their participation. The study and all the procedures were approved by the Western University Health Sciences Research Ethics Board (HSREB) team.

### 2.2 Procedure

Participants viewed rapid series of natural object images and were instructed to identify two face target images embedded in the stream. Following the stream, participants were presented with two consecutive questions to choose the presented T1 and T2 from four face image options by pressing a button. Prior to the experiment, participants viewed all target face images, each for 1 second, and they did a practice run to familiarize themselves with the task.

### 2.3 Experimental Design

We selected target images from a set of 16 face images (8 women and 8 men) sourced from the Flickr Face HQ dataset (Hebart et al., 2019). Distractors were drawn from a pool of 220 object images sourced from the THINGS object concept and object image database (Karras et al., 2019). All face images displayed smiling facial expressions and a forward gaze, but they were diverse in ethnicity. We chose faces as targets so that the target images are sufficiently distinct for participants to identify amidst the rapid stream of natural object images. Each image was converted to grayscale and resized to 220*220 pixels using MATLAB functions. Luminance matching was applied to all distractor images using the SHINE MATLAB toolbox (Willenbockel et al., 2010). The stimuli were presented using MATLAB Psychtoolbox (Pelli and Vision, 1997; Kleiner et al., 2007) on a 30-inch Dell monitor with 1200*1200 pixels resolution and a refresh rate of 59.95 Hz. Participants were seated 70 cm away from the monitor, and the stimuli covered 5 degrees of visual angle centrally on a gray background on the screen.

Each trial began with a random 600-800 ms fixation (uniformly distributed) followed by a rapid series of 15 images (13 distractors and 2 targets) each presented for 84 ms. The first target was randomly presented as either the fourth or sixth item in the stream. The second target was randomly third or seventh item after the first target, resulting in two different lag conditions, lag 3 (short lag condition) and lag 7 (long lag condition). There were at least two items that followed T2 in order to mask the second target. At the end of each trial, after a random delay of 600-800 ms (uniformly distributed), participants identified T1 and T2 respectively among 4 options with a button press (Figure 1A).

**Figure 1.**
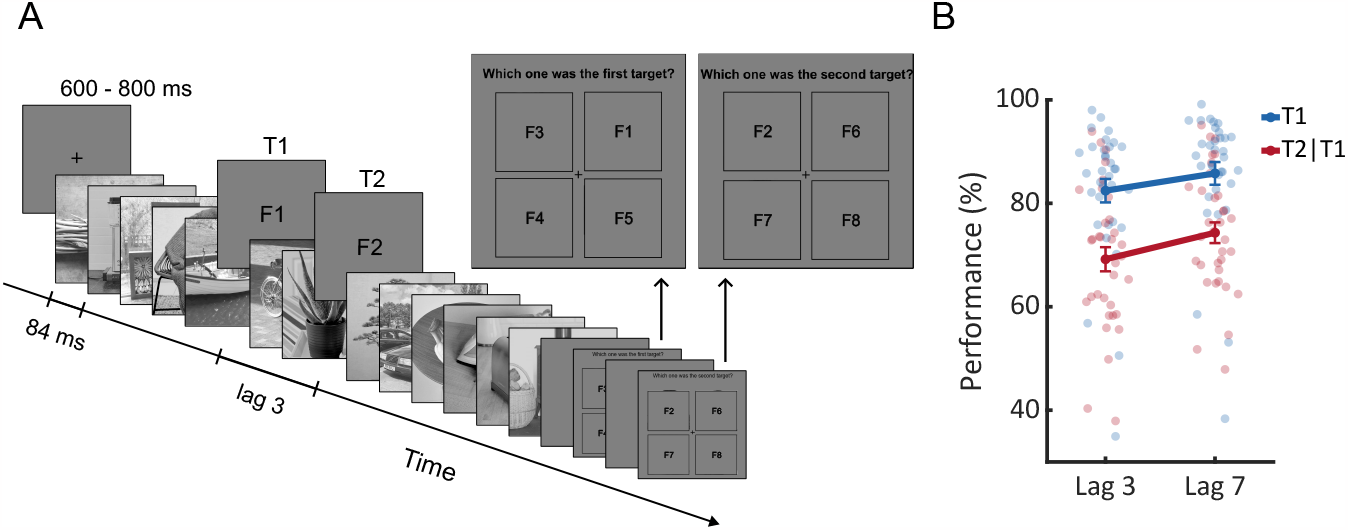
Experiment design and behavioral results. **(A)** Each trial started with a fixation for a random duration of 600 - 800 ms followed by 15 items: 2 targets and 13 distractors. Target images were randomly chosen from 16 face images (8 men and 8 women) and distractor images were randomly chosen from 220 natural object images. No image (target or distractor) was presented more than once in a trial. T1 was presented randomly as item 4 or item 6 in the stream and T2 was randomly presented at lag 3 or lag 7, each in half of the trials. At the end of each trial, after a random delay of 600 - 800 ms, participants were asked to report T1 and T2 respectively between four options by pressing a button. Face images are replaced with F1-F8 due to submission requirements. **(B)** Accuracy of reporting targets in the two lag conditions. T1 reporting accuracy is shown in blue, and T2|T1 (T2 given T1 identified correctly) reporting accuracy is shown in red. Circles show each participant’s reporting accuracy. Error bars indicate standard error (SE). There was a significant effect of lag in T1 accuracy (N = 34 participants, two-sided paired-sample t-test, lag 3 accuracy: mean = 82.44%, SE = 2.27%; lag 7 accuracy: mean = 85.78%, SE = 2.20%; t(33) = 6.84, p = 8.5e^-8^) and T2|T1 accuracy (N = 34 participants, two-sided paired-sample t-test lag 3 accuracy: mean = 69.19%, SE = 2.33%; lag 7 accuracy: mean = 74.31%, SE = 2.00%; t(33) = 5.01, p = 1.8e^-5^).

Each participant completed 22 runs, each with 32 trials for a total number of 704 trials. In each run, all 16 target images were randomly presented as T1 and T2, without repeating the same image for both targets. The same pair of images were used in both lag 3 and lag 7 conditions.

### 2.4 EEG recordings and preprocessing

Continuous EEG was recorded in a sound attenuated dimly lit room using a 64-channel Biosemi Active-Two system fitted with active-amplified electrodes. Electrodes were arranged according to the standard 10–10 system. Data were recorded at a sampling rate of 2048 Hz and all electrode impedances were kept below 20 kΩ.

EEG data preprocessing was performed using the Brainstorm MATLAB toolbox (Tadel et al., 2011). EEG data was filtered with a 0.5-30 Hz band-pass filter and downsampled to a sampling rate of 512 Hz. Bad channels were removed if their power spectral density was completely different compared to other channels. Eyeblink and eye movement artifacts were corrected by Picard ICA (Makeig et al., 1996). The data was segmented into two groups of trials with respect to the onset of T1 and T2 (−200 ms to 1000 ms) with 200 ms pre-stimulus baseline correction. Trials containing excessive movement artifacts were rejected manually, and trials with incorrect T1 responses were excluded from further analyses.

### 2.5 Multivariate pattern analysis

For each target in each lag condition, trials were labeled by their target face images. There were a maximum of M = 22 repetitions for each of the 16 target faces. Using 64 EEG channels, each time point t (from -200 to 1000 ms with 512 Hz sampling rate) in each trial included a pattern vector of 64 measurements. For each time point, we performed pairwise classification using a linear support vector machine (SVM) classifier on EEG pattern vectors (Chang and Lin, 2011). To improve the signal-to-noise ratio, M pattern vectors of each label were divided into five folds, and pattern vectors within each fold were averaged, resulting in five input observations per face target image and a total of 80 (16*5) input observations for the classifier (each contained one EEG pattern vector at time point t). Using leave-one-out cross-validation method, four averaged pattern vectors were assigned as the train set, and one as the test set. The cross-validation process was repeated 100 times with random assignments of pattern vectors into averaged folds. The output pairwise classification accuracies were then averaged over all pairs to calculate each target decoding accuracy, at time point t, for each lag condition. Decoding analysis was done separately for each participant and then averaged over all participants. The averaged decoding accuracies for all participants over time were obtained to trace the neural dynamics of T1 and T2 processing in lag 3 and lag 7 in EEG data (Mohsenzadeh et al., 2018).

To look at the stability of the neural representations over time, we extended the SVM classification procedure using a temporal generalization approach (King and Dehaene, 2014; Mohsenzadeh et al., 2018; Meijs et al., 2019). We trained the SVM classifier on time point t and tested it on all time points. This yielded a temporal generalization map where each element shows the averaged decoding accuracy over all pairs of target images.

### 2.6 Correct versus incorrect T2 responses

To evaluate the difference between correct and incorrect T2 responses in the lag 3 condition, we separated all the trials into two sets of correct and incorrect responses depending on the correct identification of T2. We then conducted pairwise classification between correct and incorrect labels to calculate the decoding accuracy time series for T2 responses. Since the number of correct and incorrect trials were different, we randomly sampled N (the number of trials in the smaller set) trials from the bigger set before averaging the trials within each input fold to ensure the same number of trials in both the correct and incorrect sets. We performed a random bootstrapping with 1000 repetitions, ensuring a similar signal-to-noise ratio for all input folds. The decoding accuracy time series for correct versus incorrect T2 responses in lag 3 condition were calculated by averaging pairwise classification accuracies over bootstrap repetitions. As above, the temporal generalization map for correct versus incorrect T2 responses in the lag 3 condition was also calculated.

### 2.7 Statistical analysis

Statistical inference of MVPA analysis relied on cluster-based non-parametric statistical tests (Maris and Oostenveld, 2007). We performed permutation-based cluster-size inference for statistical assessment of decoding accuracy time series and temporal generalization maps. For the decoding analysis, we used 10000 permutations, a 0.05 cluster defining threshold, and a 0.05 significance threshold. For the temporal generalization analysis, we used 10000 permutations, a 0.01 cluster-defining threshold, and a 0.05 significance threshold.

For statistical assessment of lag 7 and lag 3 differences in decoding analysis, we performed bootstrap tests. Thirty-four participants were bootstrapped with 1000 repetitions with substitution. In Each repetition, the average decoding accuracy time series (across participants) was divided into 6 windows, each with a length of 200 ms (from -200 ms of the target onset to 1000 ms), and the decoding accuracy within each window was averaged. This yielded a distribution of 1000 samples of average decoding accuracy per target per lag condition. We chose this window length based on earlier studies which identified different stages of target processing: an early stage occurring up to 200 ms of the target onset and a later processing stage after ∼350 ms of target onset associated with conscious report (Marti and Dehaene, 2017; Weaver et al., 2019). P-values were evaluated as the proportion of bootstrap samples that were less than the 50% chance level (null hypothesis). We then subtracted lag 7 and lag 3 distributions resulting in an empirical distribution of average decoding difference per target. To assess the difference statistically, p-values were evaluated as the proportion of bootstrap samples that were less than zero. All p-values were corrected for multiple comparisons using Benjamini-Hochberg false discovery rate (FDR) correction method (Benjamini and Hochberg, 1995). The significant threshold was set at 0.05.

For statistical assessment of the difference between lag 7 and lag 3 temporal generalizations, we performed a similar approach using 200*200 ms^2^ time windows. For each time window, a 1000 sample distribution of the difference between lag 7 and lag 3 average decoding accuracy was calculated per target. Then, p-values were evaluated as the proportion of bootstrap samples that were less than zero. All p-values were corrected by false discovery rate correction at 0.05.

## 3 Results

### 3.1 Behavioral signatures of attentional blink in naturalistic paradigm

We calculated the participants’ average accuracy of reporting T1, and T2 given T1 had been reported correctly (T2|T1) for the two lag conditions (Figure 1B). We observed a significant difference in T2|T1 accuracy between the two lags using a two-tailed paired-sample t-test (N = 34 participants; lag 3 accuracy: mean = 69.19%, SE = 2.33%; lag 7 accuracy: mean = 74.31%, SE = 2.00%; t(33) = 5.01, p = 1.8e^-5^). This, in line with previous studies, indicated that AB occurred in our study (Raymond et al., 1992). Moreover, there was a significant increase in T1 accuracy in lag 7 compared to lag 3 (N = 34 participants, lag 3 accuracy: mean = 82.44%, SE = 2.27%; lag 7 accuracy: mean = 85.78%, SE = 2.20%; t(33)=6.84, p = 8.5e^-8^). This finding is similar to earlier studies highlighting that T1 reporting accuracy can vary at different lag conditions (Olivers and Nieuwenhuis, 2006).

### 3.2 Neural representations of natural target processing depend on the lag between targets

We sought to determine if the neural representations of target image processing exhibit a similar pattern as the behavioral results. To compare the neural representations of target processing in short and long lag conditions, we calculated decoding accuracy time series for T1 and T2 in lag 3 and lag 7 conditions by performing SVM pairwise classification on EEG pattern vectors of target images (See Methods section 2.5 for more information). It is worth reiterating that all trials with incorrect T1 responses were excluded prior to the multivariate pattern analysis.

In the lag 7 condition, decoding accuracy increased significantly as early as ∼102 ms after the onset of T1 and dropped to the chance level (%50) after ∼232 ms. The early rise in decoding accuracy indicated that target identity could be decoded from the EEG data, and neural responses were resolved at the level of individual faces (Mohsenzadeh et al., 2018). After 500 ms post-target onset, decoding accuracy rose again indicating that T1 processing continued for a longer period of time (as long as ∼600 ms after the T1 onset). At ∼ 674 ms (∼86 ms after the onset of T2), T2 decoding accuracy increased significantly, while T1 decoding accuracy dropped to the chance level. The early rise in T2 decoding accuracy took ∼264 ms, and then decoding accuracy decreased to the chance level. At around 1000 ms after the T1 onset, T2 decoding accuracy rose again, indicating T2 further processing similar to what we saw in T1 decoding accuracy time series (Figure 2A).

**Figure 2.**
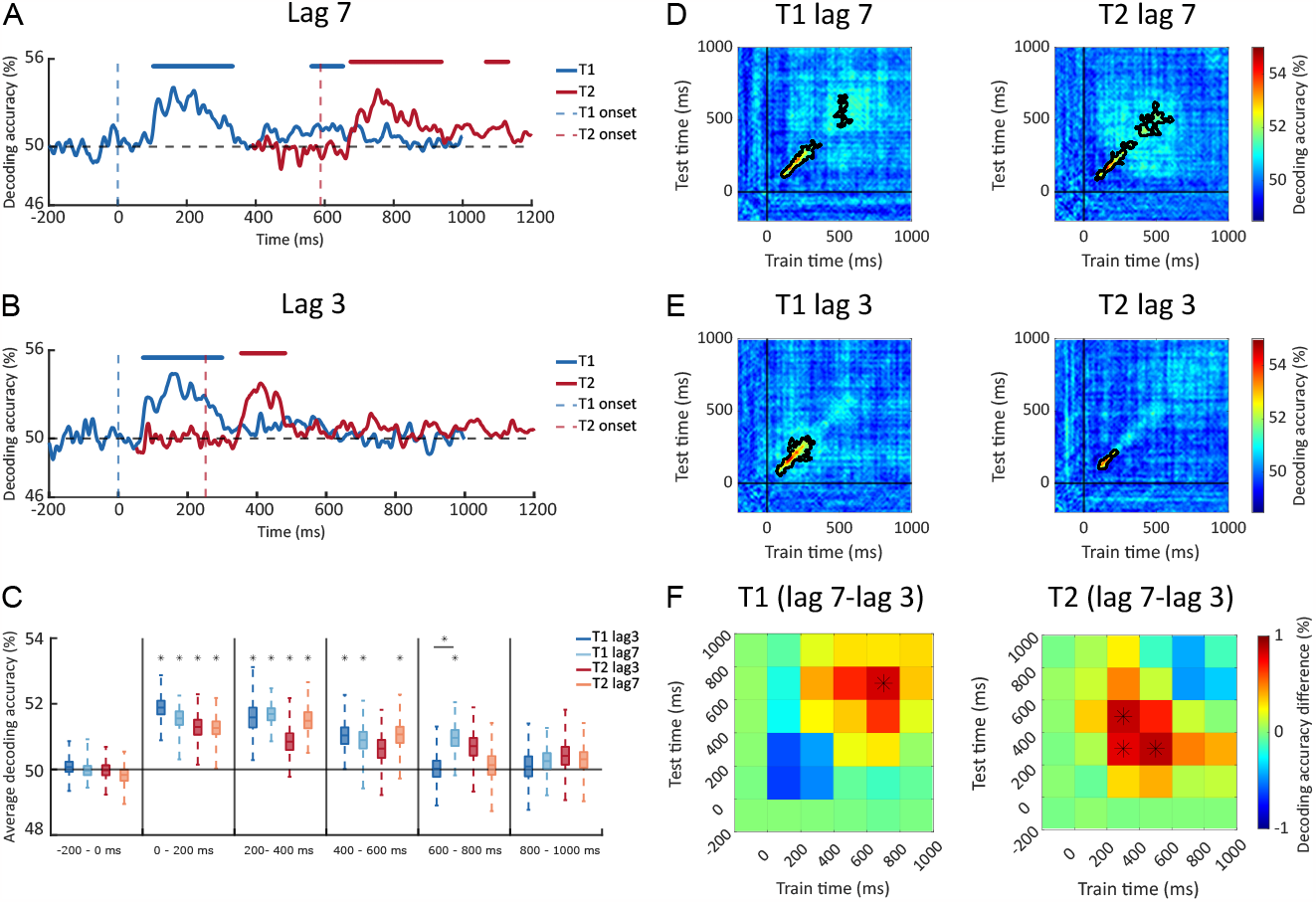
Decoding analysis. Decoding accuracy time series of T1 (blue) and T2 (red) in **(A)** lag 7 and **(B)** lag 3 conditions. The vertical dashed lines show the onset of T1 (blue) and T2 (red). The horizontal dashed line shows a 50% chance level. The lines above the graphs are significant intervals that were calculated using one-sided permutation tests and corrected with cluster correction (N = 34 participants; cluster defining threshold p < 0.05, corrected significance level p < 0.05, 10000 permutations). **(C)** The average decoding accuracy within 200 ms time windows for T1 (blue) and T2 (red) in the two lag conditions. The average decoding accuracy was calculated using bootstrap tests. The horizontal line shows a 50% chance level. The vertical lines separate time windows. Significant average decoding accuracies are shown with black stars (1000 permutation with N = 34 participants with substitution, one-sided permutation tests, false discovery rate corrected at p < 0.05). **(D)** Temporal generalization maps for T1 and T2 in lag 7 and **(E)** lag 3 condition. Warmer colors indicate higher decoding accuracy. Black vertical and horizontal lines show the corresponding target onset. Significant clusters, shown with a black contour were calculated using one-sided permutation tests and corrected with cluster correction (N = 34 participants; cluster defining threshold p < 0.01, corrected significance level p < 0.05, 10000 permutations). **(F)** Difference between lag 7 and lag 3 average decoding accuracy within 200*200 ms^2^ time windows. The average decoding accuracy was calculated using bootstrap tests. Significant time windows are shown with black stars (1000 permutation with N = 34 participants with substitution, one-sided permutation tests, false discovery rate corrected at p < 0.05).

The decoding accuracy time series of both targets appears to reflect two temporally distinct stages of processing. The early processing occurred approximately 100 to 300 ms after the target onset, while the late processing happened after 400 ms. This is consistent with previous studies suggesting that there are two stages of processing for a target: an early stage happens as early as 100-200 ms after target onset, while the second stage of processing starts later (∼400 ms) and extends for a longer time. (Marti and Dehaene, 2017; Weaver et al., 2019; Meijs et al., 2019; Alilović et al., 2021).

What we observed in the lag 3 condition was different (Figure 2B). Here, T1 decoding accuracy rose significantly at around 72 ms after the T1 onset and dropped to the chance level after ∼228 ms. T2 decoding also showed an early rise in accuracy after ∼102 ms, and it dropped to the chance level around 230 ms post-stimulus onset. The early rise in T1 and T2 decoding was similar to the early rise of target decoding in the lag 7 condition, and it reflected the discrimination of neural responses at the level of individual face images in the brain. In contrast, there was no significant time interval in either T1 or T2 decoding accuracy after 500 ms of the T1 onset. This suggests that in the lag 3 condition, the subsequent processing of T1 and T2 was inhibited, preventing both targets from the later stages of processing.

To compare lag 7 and lag 3 conditions, we directly compared the average decoding accuracy in 200 ms windows from 200 ms pre-stimulus to 1000 ms post-stimulus onset. We calculated the average decoding accuracy across all the time windows in lag 3 and lag 7 conditions for both targets by performing bootstrap tests with 1000 permutations. In each permutation, we randomly chose 34 participants with substitution, and we calculated the average decoding accuracy magnitude within each time window (for more information see Methods section 2.7). The average decoding accuracy across 1000 permutations per target per lag condition is shown in Figure 2C. We observed that decoding accuracy was at the chance level during the pre-stimulus window (−200 to 0 ms). Then decoding accuracy was significantly above the chance level in the second and third time windows (0 to 400 ms), aligning with the assumed early processing stages. There was no significant difference between lag 3 and lag 7 in either T1 or T2 average decoding accuracy in the first three windows. Later in time, in the fourth window (400 to 600 ms), we observed a chance level average decoding accuracy for T2 in lag 3 condition, confirming the suppression we observed in the decoding accuracy time series of the second target. Going forward to the fifth window (600 to 800 ms), we observed that the difference between average decoding for T1 in lag 7 and lag 3 was significant, confirming the decoding suppression in lag 3 (Figure 2B). Finally, in the last window (800 to 1000 ms) all average decoding accuracies went back to the chance level (one-sided permutation tests, false discovery rate corrected at 0.05).

Together, these results suggest that early stages of target processing (reflected by the first rise in decoding accuracy time series) were not affected much by attentional blink. However, the subsequent processing, indicated by the second rise in decoding accuracy time series, was inhibited in lag 3 compared to the lag 7 condition, where there was more temporal gap between the two targets.

### 3.3 Temporal generalization analysis confirmed the effect of lag on target representations

Temporal generalization analysis enables us to delve deeper into how visual information is processed over time and the extended patterns in temporal generalization reflect the persistent maintenance of neural representations in time (King and Dehaene, 2014; Mohsenzadeh et al., 2018). In this approach, we trained the SVM classifier on one time point and tested it on all other time points, to calculate the temporal generalization maps of targets in the two lag conditions. Similar to our observation in decoding analysis, in both lag conditions, significant decoding accuracy emerged as early as 100 ms of target onset indicating that the individual target face images were decoded using EEG data.

In lag 7 condition, we observed a narrow diagonal pattern in T1 temporal generalization map from ∼100 ms to ∼300 ms (Figure 2D). This is proposed to reflect a rapid sequence of distinct neural processes that evolved over time, which indicated early target processes (King and Dehaene, 2014; Mohsenzadeh et al., 2018). Later in time, an off-diagonal square-shaped pattern can be observed in the temporal generalization map of T1. Such a pattern would reflect that decoding was extended in time and neural representations of target processing were more sustained in this stage in contrast to the early stages where the neural representations were transient (Meijs et al., 2019). Upon examining the diagonal of T2 temporal generalization map for in lag 7 condition, a similar trend became evident. On the other hand, a very different pattern was observed in lag 3 condition. We observed a narrow diagonal pattern in temporal generalization maps of T1 and T2 early in time (100 ms to 300 ms), similar to what we found in lag 7 condition. However, the decoding pattern was smaller and more limited to the diagonal for the second target. We could not see any significant square-shaped pattern in the temporal generalization maps later in time which indicated that the later processing of T1 and T2 in lag 3 was suppressed (Figure 2E).

To compare lag 7 and lag 3, we directly compared the average decoding accuracy across 200*200 ms^2^ time windows starting from -200 ms target onset to 1000 ms by performing bootstrap tests with 1000 permutations (for more information see Methods section 2.7). Figure 2F indicates the averaged decoding difference between lag 7 and lag 3. We observed that the T1 and T2 decoding accuracy in lag 7 was significantly higher than lag 3 in later time windows (one-sided permutation tests, false discovery rate corrected at p < 0.05). Comparing lag 7 and lag 3 temporal generalizations maps of T1 and T2, we found that early processing of targets, 100-300 ms after target onset, was similar between lag 3 (AB condition) and lag 7 conditions. However, the later processing was suppressed in lag 3, but not in lag 7 as a broader decoding pattern can be seen in lag 7 condition for both targets. These findings are consistent with what we found in the previous analysis (Results section 3.2) and confirm the early target processing remained unchanged regardless of lag condition, while the higher-level processes were inhibited during attentional blink.

### 3.4 Early target representations are not affected by T2 identification

To calculate the decoding accuracy time series for T2, we used all T2 trials regardless of correct or incorrect T2 responses. This raises the question of how the neural representations differ between correct and incorrect T2 responses. To answer this question, we separated the trials based on whether T2 was reported correctly and then we calculated the decoding accuracy time series for correct versus incorrect T2 responses in the lag 3 condition, the AB condition. We observed that decoding accuracy was at the chance level until ∼500 ms after the T2 onset indicating that the early processing of correct and incorrect responses were similar (Meijs et al., 2019; Alilović et al., 2021). However, decoding accuracy of correct versus incorrect responses rose significantly approximately from 500 ms to 800 ms, showing that late stages of processing differed between correct and incorrect T2 responses (Figure 3A). Furthermore, the temporal generalization map indicated a broad square-shaped pattern around 500 to 800 ms, confirming the difference between neural representations of correct and incorrect T2 responses in higher-level processes (Figure 3B).

**Figure 3.**
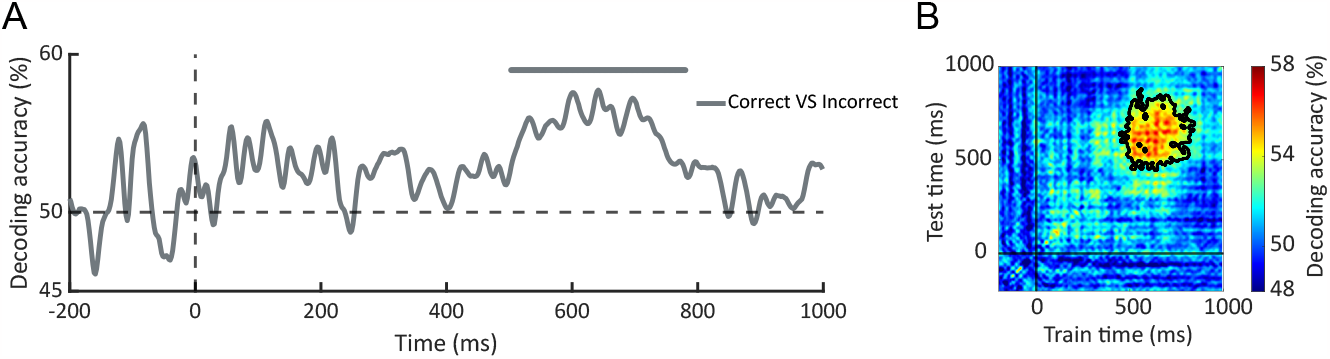
Correct versus incorrect T2 responses. **(A)** Decoding accuracy time series for correct versus incorrect T2 responses in lag 3 condition. The vertical dashed line shows T2 onset and the horizontal dashed line shows a 50% chance level. The line above the graph is significant intervals calculated using one-sided permutation tests and corrected with cluster correction (N = 34 participants, cluster defining threshold p < 0.05, corrected significance level p < 0.05, 10000 permutations). **(B)** Temporal generalization map for correct versus incorrect T2 responses in lag 3 condition. Warmer colors indicate higher decoding accuracy. Black vertical and horizontal lines show T2 onset. Significant clusters, shown with a black contour were calculated using one-sided permutation tests and corrected with cluster correction (N = 34 participants; cluster defining threshold p < 0.01, corrected significance level p < 0.05, 10000 permutations).

## 4 Discussion

The present study aimed to enhance understanding of the neural dynamics associated with target processing in a naturalistic paradigm under conditions where attentional blink does or does not occur. We used an attentional blink paradigm with two face targets and natural object distractors at two lag conditions: a short lag condition expected to induce the attentional blink, and a long lag condition where attentional blink is expected not to happen. We collected EEG and behavioral data from participants as they engaged in a task involving the identification of two face target images within a rapid stream of natural object distractors. Using multivariate pattern analysis of EEG data we calculated the neural representations of natural target processing over time under the two lag conditions.

At the behavioral level, we observed a significant impairment in the second target identification accuracy in lag 3 confirming the attentional blink phenomenon. Intriguingly, we also observed a similar pattern of decline in the identification accuracy of the first target. This finding indicated that the first target processing might be influenced by the time interval between the targets. Previously, T1 identification accuracy has been used as a baseline for T2 identification accuracy to determine whether AB occurred in a study (Chua, 2005; McLaughlin et al., 2001). However, according to MacLean and Arnell (2012), the use of T1 identification accuracy as a baseline in studies where the identifying tasks for T1 and T2 differ is not appropriate. Based on our results, even when identifying tasks for both targets are the same, T1 identification accuracy may vary across different lags. Therefore, the traditional practice of using T1 identification accuracy as a baseline to evaluate attentional blink may not be applicable in all experiments. Furthermore, these findings emphasize the importance of exercising caution when generalizing previous findings from basic stimuli to make predictions regarding behavioral performance under more naturalistic conditions (Einhäuser et al., 2007).

We presented a series of multivariate pattern analyses, in which we first found that the identity of individual faces can be decoded reliably using EEG data. It is important to note that the target stimulus set we used shared several features such as grayscale color and facial expression; therefore, the maximum decoding accuracy was relatively low (∼ %56). Moreover, comparing target representations across lag 7 and lag 3, we found that the early processing of both T1 and T2 was not different between the two lag conditions. This might suggest that attentional blink does not affect the early processing of targets. However, we observed a chance level decoding accuracy after 300 ms from the target onset in lag 3, indicating that the later processing of targets was suppressed in lag 3, reflecting attentional blink. In lag 7, on the other hand, where there was an additional time between targets, another significant rise, reflecting late processing stages could be seen in the decoding accuracy of targets, indicating that target processing was preserved for a longer duration. The similar pattern of T1 and T2 decoding accuracy suggests that both T1 and T2 late processing are inhibited when they emerge closely one after the other (AB condition).

Moreover, we confirmed the suppression of target representations beyond 300 ms post-target onset (at higher-level processing) by evaluating the temporal generalization results. The temporal generalization maps showed a narrow diagonal pattern around 100 to 300 ms after the target onset indicating that the target representations evolve transiently in the first stages of processing. Later than this, however, there was a broad square-shaped pattern in the temporal generalization of targets in the lag 7 condition, suggesting the involvement of higher-level processes in target identification. In contrast, such patterns cannot be seen in the lag 3 condition, indicating that the high-level processes were suppressed during the attentional blink. Similar to previous results, these findings imply that both T1 and T2 representations are more sustained when there is enough time between the targets. Meijs et al. (2019) suggested a similar pattern in the temporal generalization maps of T2, when T2 (letter targets) was correctly identified.

Our results are in opposition to limited-capacity theories suggesting that during a rapid presentation of images, our attention system’s resources become occupied with processing the first target when it is presented, yielding insufficient resources for processing the second target; thus attentional blink happens (Dux and Marois, 2009). In certain models within this category, it has been proposed that T2 processing should be delayed until T1 processing is completed, but in situations where there is limited time between targets (short lag conditions), T1 processing may not have sufficient time to finish before the arrival of T2; as a result, T2 cannot be adequately processed, leading to attentional blink (Chun and Potter, 1995; Marti et al., 2012). In our results; however, the early stages of target processing can be seen for the second target as well as the first target in the lag 3 condition. Furthermore, both targets’ late processing was restricted which suggests that the available resources are engaged with both targets simultaneously rather than completing the first target processing before starting the second one. On the other hand, our findings are more in line with attentional control theories (Nieuwenstein et al., 2005; Olivers and Meeter, 2008; Olivers et al., 2007). These theories posit that attentional blink occurs due to the interference of targets and distractors’ attentional sets. Attentional control theories explain why T1 representations, just as T2 representations, were suppressed in the lag 3 condition. Although our results do not directly investigate the impact of distractors on the neural representations of targets, It would be compelling for future studies to explore and compare the neural representations of both distractors and targets, as well as their potential interactions.

Next, we investigated the difference between neural representations of correct and incorrect responses of the second target. We observed no significant difference between correct and incorrect T2 responses at early times indicating that early processing of the target is intact regardless of correct identification of target (Meijs et al., 2019; Alilović et al., 2021). Nonetheless, a significant difference can be seen later in time, meaning that the late stages of processing were different when participants correctly identified the second target.

Taking these results into account, we observed two stages of processing in the neural representations of face targets in an attentional blink paradigm with natural stimuli. This is aligned with previous studies that used an attentional blink paradigm with basic stimuli (such as digits, letters, or objects without background) (Kaiser et al., 2016; Alilović et al., 2021; Meijs et al., 2019). The early stage, which appeared as a significant rise in the decoding accuracy of the targets, started as early as 100 ms and continued until 300 ms after the target onset. This early processing was similar in both T1 and T2 in lag 3 and lag 7 conditions, meaning that the early target processing was not affected by attentional blink. One possible implication could be that the early stages of target processing are processed unconsciously; thus, regardless of the lag condition or target location in a rapid series, early processing occurs similarly. In our findings, the early stage was even similar between correct and incorrect T2 responses, confirming that the immediate processes following the target onset occur unconsciously. The second stage of processing can be seen around 400 to 800 ms after the target onset only in the lag 7 condition, and there was a significant difference between neural representations in lag 7 and lag 3, showing that this stage of processing is affected by attentional blink. Therefore, the late stages of processing might be associated with the conscious identification of the target. Additionally, we observed that this late pattern in the neural representations of the target was similar for both T1 and T2, indicating that the late stages of both targets’ processing in an attentional blink paradigm depend on the time interval between the two targets.

In conclusion, first, we demonstrated that neural representations of targets in a rapid presentation of natural object images can be decoded from neural activity measured by EEG. Second, we showed that the late neural representations of targets are suppressed if the second target is presented within 200 - 500 ms of the first target, the attentional blink period; thus, both the first and the second target identification accuracy is reduced. On the other hand, the early representations of targets are similar regardless of the targets’ lag or the behavioral performance of participants. Our research extends the existing understanding of attentional blink by investigating naturalistic paradigms and employing multivariate pattern analysis. The implications of our findings are substantial for attentional blink theories, highlighting the need for their extension in order to account for naturalistic paradigms and new findings.

## Acknowledgments

This study was supported by the Canada First Research Excellence Fund (CFREF) through a BrainsCAN grant to Y.M., and a Vector Institute Research Grant to Y.M. Also, M.J. received a graduate student scholarship from CFREF Brain-sCAN. The authors would like to thank Diana Dima and Mehrdad Kashefi for helpful comments on the manuscript and Ali Tafakkor and Saba Charmi Motlagh for helpful discussions.

